# The left posterior angular gyrus is engaged by autobiographical recall not object-semantics, or event-semantics: Evidence from contrastive propositional speech production

**DOI:** 10.1101/2022.04.04.487000

**Authors:** Gina F. Humphreys, D. Halai Ajay, M. Branzi Francesca, Matthew A. Lambon Ralph

## Abstract

The angular gyrus (AG) has been implicated in a myriad of cognitive functions. Using the previously under-studied naturalistic task of propositional speech production, we investigated the engagement of the left posterior AG (pAG) by three forms of memory: 1) episodic/autobiographical memory, 2) object semantic-memory, and 3) event-semantic knowledge. We conducted an ALE meta-analysis, followed by an fMRI study. The meta-analysis showed that pAG is only engaged as part of the propositional speech network when the task carries an autobiographical component. This finding was supported by the fMRI results, which also showed that: 1) pAG was positively engaged during autobiographical memory retrieval; 2) pAG was strongly deactivated for definitions of object semantics and non-propositional speech; 3) pAG activation increased with the degree to which the event descriptions relied on autobiographical information; 4) critically, the pAG showed a different pattern to known semantic representation regions (e.g., ventral anterior temporal lobe (vATL)) thereby providing clear evidence that the pAG is not acting as a semantic hub. Instead, the pAG activation profile directly mirrored that found in the wider autobiographical retrieval network. We propose that information stored elsewhere in the episodic system is temporally buffered online in the pAG during autobiographical retrieval/memory construction.

## Introduction

A long history of neuropsychology and neuroimaging has implicated the lateral parietal cortex (LPC) in a myriad of cognitive functions. Whilst highly sophisticated models exist for each domain in isolation, the majority of theories fail to account for the variety and potential functional overlap of LPC activities. There are two alternative ways that the LPC might be organised: 1) A form of ‘neuromarquetry’ in which LPC contains numerous discrete sub-regions each recruited by different tasks and serve distinct underlying cognitive functions; or 2) shared computations - multiple cognitive activities rely, in common, upon a small number of common underlying computations that arise from the LPC and its patterns of connectivity to the wider neural network. Resolving this puzzle will necessitate direct cross-domain comparisons. Two domains that have heavily implicated the LPC, specifically the left PGp subregion of the angular gyrus (hereon pAG), are episodic memory and semantic memory retrieval. For instance, one model proposes that the pAG forms an episodic buffer for retrieved episodic information (Vilberg & Rugg, 2008; A. D. Wagner, Shannon, Kahn, & Buckner, 2005). An alternative to the episodic hypothesis is that pAG is a semantic-hub that stores multi-modal semantic information (Binder, Desai, Graves, & Conant, 2009; Geschwind, 1972; Geschwind, Quadfasel, & Segarra, 1968). A related though more specific alternative to the semantic-hub hypothesis is that the pAG stores “semantic event” information per se, whereas “object” information is stored elsewhere in the semantic system, such as the ventral anterior temporal lobe (vATL) (Binder & Desai, 2011). Indeed, centred on the anterior fusiform gyrus, the ATL is widely regarded as a key hub for semantic representation (Lambon Ralph et al., 2017). Alternatively, others have proposed domain general models, for instance the theory that pAG is implicated by any “internally-generated” cognition (including semantic memory or episodic memory retrieval, as well as other processes) (Andrews-Hanna, 2012; Buckner, Andrews-Hanna, & Schacter, 2008), or that it is a part of a broader IPL multi-modal sequential buffer (Humphreys, Jackson, & Lambon Ralph, 2020; Humphreys & Lambon Ralph, 2015; Humphreys, Lambon Ralph, & Simons, 2021). In addition to the confusing number of potential roles of the AG, the majority of the studies have investigated “input” tasks (e.g., making a decision on written/auditory words or pictures), whereas relatively few have looked at “output” tasks, such as speech production.

### Direct within-study comparisons

A test of these alternative hypotheses of pAG function requires direct cross-domain within-study comparisons. A recent functional magnetic resonance imaging (fMRI) study manipulated internally vs. externally directed attention, and episodic retrieval vs. semantic retrieval (Humphreys, Jung, & Lambon Ralph, 2022). There were four conditions, two involving internally directed attention (semantic retrieval or episodic retrieval) and two involving externally directed visual attention (real-world object decision, or scrambled pattern decision). The pAG, a region that overlaps with the default mode network (DMN), was found to show strong positive activation during episodic retrieval, with the level of activation correlating with item-specific memory vividness ratings. In contrast, in the semantic retrieval task and the object decision task the pAG showed deactivation relative to rest and relative to the control task, with the level of deactivation correlating with task difficulty (stronger deactivation for the harder items). Functional and structural connectivity analyses found that the same pAG region of interest (ROI) showed connectivity with the wider episodic retrieval network, including the medial temporal lobes, as well as medial prefrontal, precuneus, and posterior cingulate cortex (Buckner & Carroll, 2007; Daselaar et al., 2009; Ranganath & Ritchey, 2012), consistent with existing evidence (Caspers et al., 2008; Caspers et al., 2011; Cloutman, Binney, Morris, Parker, & Lambon Ralph, 2013; Humphreys et al., 2022; Uddin et al., 2010). Combined, these data suggest functional involvement of the pAG in episodic rather than semantic retrieval or the processing of all forms of internally-generated thought. Nevertheless it does not preclude the possibility that the central function of pAG is “semantic event” knowledge since episodic retrieval might necessitate the activation of event semantics. The results are also consistent with the Parietal Unified Connectivity-biased Computation (PUCC) model (Humphreys et al., 2020; Humphreys & Lambon Ralph, 2015; Humphreys et al., 2021), whereby the emergent expressed function will depend on what sources of information and influences arrive at each subregion; the pAG connects with the episodic retrieval network and hence is more engaged by tasks that involve episodic retrieval (Humphreys et al., 2020; Humphreys et al., 2022; Rugg & Vilberg, 2013; Sestieri, Corbetta, Romani, & Shulman, 2011). Note that PUCC and the “episodic buffer” model are not mutually exclusive, PUCC simply offers a generalised mechanistic explanation as to how an episodic buffer function might emerge.

### Propositional speech production as a useful paradigm to investigate pAG function

The current study used a propositional speech production paradigm to determine the underlying function of (left) pAG whilst also addressing a number of existing limitations in the literature. Speech production was used for four primary reasons: 1) relatively little is known about the involvement of IPL, including AG, in expressive tasks since existing theories are based almost exclusively on “input” tasks (e.g., word recognition); 2) Propositional speech is inherently “internally-generated” and thus makes the ideal testing environment to determine to what extent the pAG is recruited for a variety of internally-generated domains (semantic, episodic etc.); 3) Speech production can be a powerful and flexible paradigm to investigate cognitive function as one can carefully manipulate the speech topic (e.g., semantic definition vs. autobiographical description) and thereby influence the degree to which different subsystems are drawn upon to produce speech content; 4) Finally, narrative speech production is a time-extended behaviour that therefore provides insights into the internal operation of a “temporal buffer” model. The current literature on speech production and the pAG is sparse, and provides mixed support for its engagement. A few studies have found that pAG is actively engaged when participants answer questions, such as “tell me about the last family gathering you attended” (Braun et al., 1997; Brownsett & Wise, 2009). However, others have found greater pAG activation when performing non-verbal tongue movements compared to meaningful speech production (providing definitions of a noun) (Geranmayeh et al., 2012). The reason for these discrepant results is unclear. For instance, of relevance here, the studies varied in terms of speech topic; LPC activation was observed for the autobiographical description task but not a noun-definition task. However, the existing research is hard to interpret because there are numerous other variations across studies (e.g., speech duration, PET vs. fMRI, covert vs. overt speech, variety of tasks utilised).

The current study examined whether the (left) pAG is engaged during speech production, and if so, what cognitive variables modulate pAG engagement. We first conducted a meta-analysis of the existing propositional speech production neuroimaging studies in order to determine the reliability of pAG effects, and the extent to which they are influenced by task-related variables. Whilst meta-analyses are powerful techniques that highlight generalizable results across studies (and across variations in methods, analyses, etc.), they can blur data thereby losing the anatomical precision that can be derived from direct within-participant manipulations. Therefore, we conducted a within-subject cross-domain fMRI study in which we directly manipulated narrative speech content. Specifically, the participants provided verbal responses to a number of questions during the fMRI task. There were four varieties of question: 1) autobiographical condition: the participants answered questions asking about specific past events (e.g., describe the last time you drank a cup of tea); 2) semantic-event condition: the participants described how a particular event would typically unfold (e.g., describe how you would typically make a cup of tea); 3) semantic-object condition: the participants provided semantic-definitions of objects (e.g., what is tea?); and, 4) control condition: a non-semantic speech control task (recite the months of the year).

## Methods

### Meta-analysis

#### Study selection

The meta-analysis comprised 25 PET and fMRI studies that included a sentence- or narrative-level propositional speech production paradigm compared to a control condition or resting baseline (see Supplementary Table S1 for a list of included studies). Our inclusion criteria were <u>any</u> fMRI/PET study that had used a sentence- and/or narrative-level speech production task, and reported peak activation in standard space (Talaraich or MNI) based on whole-brain statistical comparisons. Studies were primarily selected from existing comprehensive meta-analyses that investigated speech production or autobiographical memory retrieval, provided that the studies met our inclusion criteria (Kim, 2013; Spreng, Mar, & Kim, 2009; Walenski, Europa, Caplan, & Thompson, 2019), and supplemented by screening for studies that met the inclusion criteria in Google Scholar searches (using keywords: “speech production”, “speech generation”, “propositional speech”, “sentence”, “narrative”, “verbal description”, AND “fMRI”, “PET”), and screening of citations in selected papers. Examples of typical contrasts were: define a noun > counting, sentence generation (based on key words) > sentence reading, sentence generation (based on picture) > rest. Amongst the selected papers, six studies included autobiographical manipulations, such as tasks involving self-referential narratives recounting autobiographical memories and personal stories (e.g., the participants were asked questions such as “tell me about the last family gathering you attended” (Brownsett & Wise, 2010), or “recount a recent event from memory” (Braun et al., 1997; Braun et al., 2001) or “describe where you lived as a child” (Blank, Scott, Murphy, Warburton, & Wise, 2002) (see Table S1 for a full list of included studies). This is in contrast to tasks without an autobiographical component (e.g. “define an apple” (Geranmayeh, Wise, Mehta, & Leech, 2014), or “generate a sentence using the words: throw ball child” (Haller, Radue, Erb, Grodd, & Kircher (2005)).

#### ALE analyses

The ALE analyses were carried out using GingerAle 3.0.2 (Eickhoff et al., 2009; Laird et al., 2005). All activation peaks were converted to MNI standard space using the built-in Talaraich to MNI (SPM) GingerALE toolbox. Analyses were performed with voxel-level thresholding at a p-value of 0.001 and cluster-level FWE-correction with a p-value of 0.05 over 10,000 permutations. In order to identify the network engaged by propositional speech production, the initial analysis was performed including all 25 studies. Then, secondary analyses were conducted in order to examine specific task-related effects. In particular, we sought to determine the extent to which the activation likelihood maps were influenced by the inclusion of studies using autobiographical content. This was done by removing the six autobiographical studies and re-running the ALE analysis using the remaining 19 studies. Since changes in the activation likelihood maps from the first analysis (25 studies) with the second analysis (19 studies) could be due to reduction in power rather than an influence of autobiographical task per se, we performed additional analyses in which six studies from the original pool were excluded at random, the results of which could be compared to the new 19 study-pool. The random study-exclusion analysis was performed 10 times in order to observe which clusters from the activation likelihood maps were stable, thereby implying they form a core feature of the propositional speech network, and which clusters varied, thereby implying task-modulation.

### fMRI study

#### Participants

Twenty-eight participants took part in the fMRI study (average age = 27.91 years, SD = 6.19; N female = 20). All participants were native English speakers with no history of neurological or psychiatric disorders and normal or corrected-to-normal vision. The fMRI experiment was approved by the local ethics committee. Written informed consent was obtained from all participants.

#### Task design and procedures

There were four experimental conditions, with 36 items per condition: autobiographical description, semantic-event description, semantic object description, and control condition. Each trial involved two phases; memory retrieval phase and response phase. In the memory retrieval phase a question was visually presented on the screen for 5.5 seconds during which the participants could read the question and prepare their response. There was then a brief fixation cross (jittered from 0-100 ms), and then a green X that signalled the beginning of the response phase. In the response phase, the participants were asked to answer the question aloud and the responses were recorded using a MR-compatible microphone. The subjects were given 14 seconds to respond and were encouraged to speak for the full-duration. The trials were presented using an event-related design with the most efficient ordering of events determined using Optseq (http://www.freesurfer.net/optseq) with the constraint that within each run the total time spent in “rest” matched to the total duration of each condition. Null time was intermixed between trials and varied between 0 and 15 seconds (average = 3.82 seconds, SD = 5.95) during which a fixation cross was presented. The experiment was presented in 3 runs which included 12 items from each condition. Each run lasted 446 seconds, and the order of runs was counterbalanced across participants. The content of each question was matched across conditions, but the same person never saw more than one version of each item. See supplementary Table S2 for a complete list of experimental items.

#### Autobiographical condition

Here the participants were presented with a variety of questions probing autobiographical events (e.g., describe the last time you had a cup of tea). Participants were instructed to describe the event using as much highly specific spatio-temporal contextual detail as possible. For example, where the event took place, who they were with, when did it happen, or any further sites/scenes/sounds associated with the event.

#### Semantic-event condition

These questions had a similar content to the autobiographical condition but were phrased to refer to a generalised semantic-event rather than a personal autobiographical experience (e.g. describe how you would typically make a cup of tea).

#### Semantic-object description

These questions were again of a similar content but instead involved an object-description rather than an event (e.g. what is tea?)

#### Control condition

The control condition was “recite the months of the year”. The participants continued the task for the full 14 seconds (i.e., they might complete more than one full cycle of the year). This task was used as a control for any activation associated with speech apart from that association with propositional narratives.

After the fMRI session, the participants were asked to rate each semantic-event question on a 1-5 scale in terms of how vividly they could picture the event (vividness rating), how frequently they experienced the event (event novelty rating), the extent to which they answered the question based on general knowledge of how the event typically unfolds vs. a highly specific memory of experiencing the event (general-knowledge vs. episodic-memory strategy), and how difficult they found the question to answer (difficulty rating).

### fMRI acquisition

MRI data were collected using a Siemens 3 T PRISMA system. T1-weighted images were acquired using a 3D Magnetization Prepared RApid Gradient Echo (MPRAGE) sequence [repetition time (TR) = 2250ms; echo time (TE) = 3.02 ms; inversion time (TI) = 900 ms; 230 Hz per pixel; flip angle = 9°; field of view (FOV) 256 × 256 × 192 mm; GRAPPA acceleration factor 2]. Functional data were acquired using a multi-echo multi-band (MEMB) blood oxygenation level dependent (BOLD)-weighted echo-planar imaging (EPI) pulse sequence. The MEMB sequence had the following parameters: (TR = 1792 ms; TEs = 13, 25.85, 38.7, and 51.55 ms; flip angle = 75°; FOV = 192 mm × 192 mm, MB factor = 2, in-plane acceleration = 3, partial Fourier = 7/8). Each EPI volume consisted of 46 axial slices in descending order (3 x 3mm) covering the whole brain (FOV = 240 x 240 x 138 mm). For each session, 252 volumes were acquired.

### Preprocessing

All raw DICOM data were converted to nifti format using dcm2niix. The T1 data were processed using FSL (v5.0.11) (Jenkinson, Beckmann, Behrens, Woolrich, & Smith, 2012; Smith et al., 2004; Woolrich et al., 2009) and submitted to the ‘fsl_anat’ function. This tool provides a general processing pipeline for anatomical images and involves the following steps (in order): 1) reorient images to standard space (‘fslreorient2std’), 2) automatically crop image (‘robustfov’), 3) bias-field correction (‘fast’), 4) registration to MNI space (‘flirt’ then ‘fnirt’), 5) brain extraction (using fnirt warps) and 6) tissue-type segmentation (‘fast’). All images warped to MNI space were visually inspected for accuracy. The functional MEMB data were pre-processed using a combination of tools in FSL, AFNI (v18.3.03) (Cox, 1996) and a python package to perform TE-dependent analysis (DuPre et al., 2020; Kundu et al., 2013; Kundu, Inati, Evans, Luh, & Bandettini, 2012). Despiked (3dDespike), slice time corrected (3dTshift, to the middle slice), and realigned (3dvolreg) images were submitted to the “tedana” toolbox (max iterations = 100, max restarts = 10), which takes the time series from all the collected TEs, decomposes the resulting data into components that can be classified as BOLD or non-BOLD based on their TE-dependence, and then combines the echoes for each component, weighted by the estimated T2* in each voxel, and projects the noise components from the data (Kundu et al., 2012; Poser, Versluis, Hoogduin, & Norris, 2006). The resulting denoised images were then averaged, and the mean image coregistered to T1 (flirt), warped to MNI space (using fnirt warps and flirt transform), and smoothed with 6 mm FHWM Gaussian kernel.

#### General Linear Modelling (GLM)

The data were analysed using a general linear model using SPM12. At the individual subject level, each condition was modelled with a separate regressor and events were convolved with the canonical hemodynamic response function. At the individual level, each task condition was modelled separately against rest. Group analyses were conducted using standard voxel height threshold p < .001, cluster corrected using FWE p < .05. We have argued elsewhere that a comparison of a task > rest is an important and informative contrast (Humphreys, Hoffman, Visser, Binney, & Lambon Ralph, 2015; Humphreys et al., 2022; Humphreys & Lambon Ralph, 2015). Given the involvement of pAG in the DMN, it is of critical importance to consider whether a task positively or negatively engages the region relative to rest. Whilst many tasks generate deactivation in the AG, this is not always the case and the handful of task types that do positively engage the pAG might be crucial sources of evidence about its true function (Humphreys et al., 2022). Contrasts between a cognitive task of interest vs. an active control condition are ambiguous because the difference could result from 1) greater positive activation for the task or 2) greater *deactivation* for the control. This issue becomes even more important when considering the impact of task difficulty on activation and deactivation in this region. A straightforward expectation applied to almost all brain regions is that if a task critically requires the pAG then the pAG should be strongly engaged by that task. Indeed, this is the pattern observed in the vATL where semantic tasks are known to positively engage the vATL relative to rest, whereas non-semantic tasks either do not modulate or deactivate vATL (Humphreys et al., 2015). Perhaps one of the major motivations for considering task (de)activation relative to ‘rest’ is that ‘rest’ can be used as a common constant reference point across tasks. This is particularly important when conducting cross-domain comparisons. For instance, when one is examining a single cognitive domain it is possible to use a domain-specific baseline, i.e., by contrasting a task that places strong demands on the particular cognitive system vs. a task with lower demands (e.g. remember > know in an episodic memory tasks, or words > non-words in a semantic memory task). Since the same is not possible across cognitive domains, rest acts a common constant for cross-domain comparisons, even if the true cognitive interpretation of ‘rest’ is unclear. Nevertheless, for completeness, analyses were replicated by contrasting each experimental task relative to the control task and importantly the results remain largely unchanged, as reported in the text and shown in Supplementary Fig S3.

#### ROI Analyses

Targeted ROI analyses were performed using a left pAG ROI. The pAG ROI composed of a 8mm sphere that was anatomically located in the PGp subregion (MNI coordinates: - 48 -64 34) in a region overlapping the DMN. This specific subregion of the pAG was chosen since it was identified from a meta-analysis on the area to show maximal likelihood of activation across semantic and episodic studies in large-scale meta-analysis (Humphreys & Lambon Ralph, 2015), thereby making it an ideal location to test the experimental manipulation of episodic and semantic-memory retrieval systems. The results from the pAG were contrasted to those from the semantic hub region within the left vATL (MNI coordinates -42 -24 -27) in a region identified from the semantic > non-semantic contrast in a study using 69 participants (Humphreys et al., 2015)), as well as the left hemisphere autobiographical network which included the perirhinal cortex, posterior cingulate cortex/precuneus, medial prefrontal cortex, and lateral middle temporal gyrus (as identified in a meta-analysis of autobiographical studies (Spreng et al., 2009)). This latter network is sometimes referred to as the posterior medial episodic system and overlaps substantially with the DMN (Buckner et al., 2008; Ranganath & Ritchey, 2012). If the pAG were a semantic hub then one would predict a similar response to the semantic network. In contrast, if the pAG were primarily involved episodic/autobiographical processes, one would expect a similar pAG response to the wider autobiographical retrieval network.

#### Regression Analyses

To foreshadow the results, the pAG was found to be strongly engaged by the autobiographical task, but also to a minimal extent by the event-semantic task (and deactivated by the other two tasks). We predicted that the level of event-semantic activation would be modulated by the memory retrieval strategy used by participants to describe each event: the extent to which the participant draws on the semantic or autobiographical memory systems. To test this hypothesis, a parametric modulator was added to the semantic-event condition to examine the relationship between certain variables and neural activation. These variables were derived from the item-ratings provided by the participants. There were three rating scales – event novelty, event vividness, and the extent to which the participant based their response on general-knowledge vs. episodic memory (hereon referred to as episodic memory strategy). In addition to these rating variables, we also included an additional variable computed based on the participants’ verbal responses - response similarity analysis - as detailed below. (Note that it was not possible to include the difficulty ratings as a regressor due to the exceedingly low variation in participant rating scores (on the 1-5 difficulty scale the most common response was “1” (very easy), with a mean rating of 1.83, SD = .94)).

1. *Event novelty*: how frequently do you experience each event? This factor might influence the reliance on a semantic vs. autobiographical memory retrieval strategy since highly frequent events would be expected to be more semanticised (i.e., will have developed an “event-schema”) whereas a participant might base their answer for novel events on a more highly specific, episodic memory.
2. *Event vividness*: how vividly can you remember last experiencing each event? More vivid memories might include more autobiographical spatio-temporal contextual details.
3. *Episodic memory strategy*: to what extent would you base your response on general knowledge of how the event typically unfolds vs. a highly specific memory of experiencing the event? An event based on general knowledge would by definition recruit semantic memory, whereas a highly-specific memory would be more episodic in nature.
4. *Response similarity analysis*: Finally, we also coded the verbal in-scanner responses from the fMRI participants. Semanticised events are long-term representations that, through repetition and rehearsal, become more stereotyped, general and independent of any one episode. Accordingly, they are reflected in consistent generalised productions across participants. Latent Semantic Analysis (LSA) (Landauer & Dumais, 1997) is methodology that can be used to measure the semantic similarity across items. LSA utilises a large corpus of several millions words to calculate a co-occurrence matrix, registering which words appear in similar contexts. It operates on the principle that words that frequently co-occur in similar contexts tend to be more semantically related. Here LSA was performed on the fMRI verbal responses in order to code the level of semantic similarity across responses for each question. Through the utilisation of a large corpus of natural language, LSA provides the user with vector-based representations of the meanings of words, which when linearly combined across words can represent the meanings of full narratives (Hoffman, Loginova, & Russell, 2018). Specifically here, for each prompt, we obtained an LSA value reflecting the associative strength from each response to every other response using a cosine similarity metric. These were averaged to give a composite LSA score for each prompt. Analyses were performed using the LSA and SnowballC packages in R (see: https://cran.r-project.org/web/views/NaturalLanguageProcessing.html) (Hoffman et al., 2018). Items with high LSA similarity scores could be regarded as more semantically similar (which would be expected for stereotyped, general semanticised memories) than those with lower scores (which would be expected for a collection of individualised episodes). The average LSA score for each item was added as a regressor to the GLM analysis

## Results

### Meta-analysis

The results from the primary ALE analysis of 25 propositional speech production studies revealed a fronto-temporo-parietal network. In the fronto-temporal network, significant clusters were observed in left precentral gyrus, inferior frontal gyrus (IFG, BA44), supplementary motor area, and middle temporal gyrus. In parietal cortex, separate clusters were observed in dorsal LPC (IPS/SPL), ventral LPC (angular gyrus), and precuneus (see Figure 1 and Table S4).

**Figure 1:**
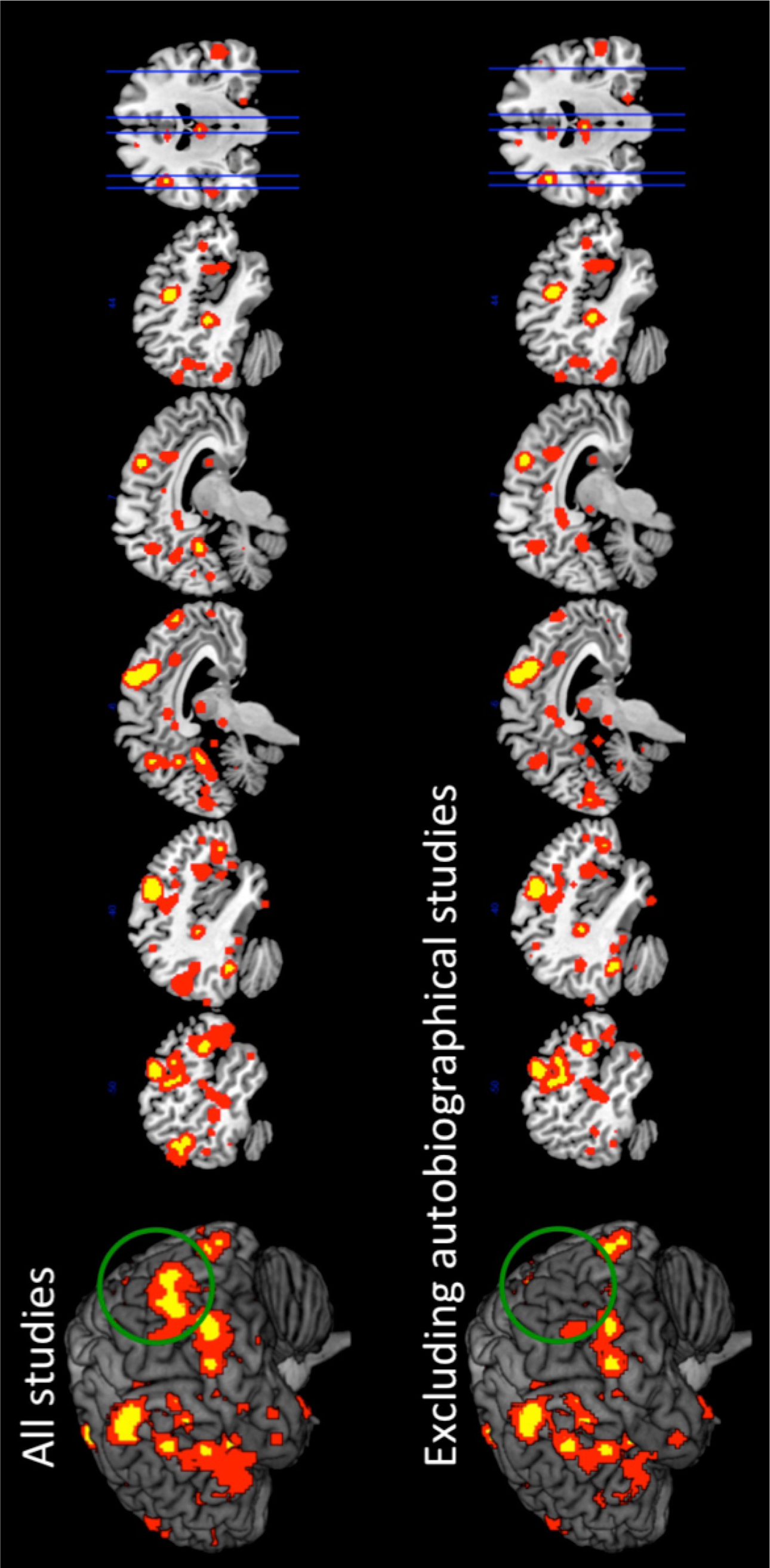
*Top panel*. The results from the ALE meta-analysis when including all 25 propositional production studies. *Bottom panel*. The results from the ALE meta-analysis after excluding autobiographical studies. Yellow indicates clusters significant at p < 0.001 (with cluster-level FWE-correction of p < 0.05), and red indicates clusters corrected at the lenient p < .05 threshold.

Secondary analyses were performed in order to determine the stability of the network across domains. Specifically, we sought to determine which clusters form core parts of the propositional speech network that are unaffected by the nature of the task, and which clusters vary depending on task domain, such as whether the task involves autobiographical memory retrieval. When the six autobiographical tasks were dropped from the analysis the fronto-temporal and dorsal parietal peaks remained constant, whereas the ventral parietal cortex (AG and precuneus) were no longer reliably activated, even when the analysis was performed at a less stringent statistical threshold (uncorrected, p < .05) (see Figure 1). The likelihood that the modulation of ventral parietal cortex was influenced by task-domain, rather than a reduction in power, was supported by the series of random study-exclusion analyses. Here we found that random deselection of six studies produced the same results as the primary ALE analysis; the pAG/precuneus was only reliably activated in the cases whereby the autobiographical studies were included, and upon inspection the AG/precuneus clusters were driven solely by those studies. That is, when the pAG and precuneus clusters were present in the ALE results, they were solely driven by peaks from the autobiographical studies. Taken together, these analyses suggest that the pAG (and precuneus) involvement is not general to propositional speech but rather to autobiographical retrieval (see Table 1). Together these data suggest that pAG does not form part of the core propositional speech network, rather it is only reliably activated when a task places requirements on the autobiographical memory system.

**Table 1.** The neural areas that showed significant clusters from the ALE analysis when including 1) all 25 studies, 2) Excluding autobiographical studies, 3) The random study-exclusion analyses, version1 (V1) to version 10 (V10). + indicates that the area was likely to show activation, with – denoting the absence of activation evidence.

|  |  | All studies | Excluding Autobiographical studies | V 1 | V 2 | V 3 | V 4 | V 5 | V 6 | V 7 | V 8 | V 9 | V 10 |
| --- | --- | --- | --- | --- | --- | --- | --- | --- | --- | --- | --- | --- | --- |
| N studies | Cluster | 25 | 19 | 19 | 19 | 19 | 19 | 19 | 19 | 19 | 19 | 19 | 19 |
| N autobiographical studies |  | 6 | 0 | 5 | 3 | 3 | 5 | 4 | 4 | 5 | 5 | 3 | 5 |
| Parietal | IPS/SPL | + | + | + | + | + | + | + | + | + | + | + | + |
|  | pAG cluster 1 | + | - | + | - | - | + | + | + | + | + | + | + |
|  | pAG cluster 2 | + | - | + | + | + | + | + | + | + | + | + | + |
|  | Precuneus cluster 1 | + | - | + | - | - | + | + | + | - | + | - | + |
|  | Precuneus cluster 2 | + | - | + | - | - | + | + | + | - | + | - | - |
| Temporal | MTG | + | + | + | + | + | + | + | + | + | + | + | + |
| Frontal | SMA | + | + | + | + | + | + | + | + | + | + | + | + |
|  | Precentral gyrus | + | + | + | + | + | + | + | + | + | + | + | + |
|  | IFG BA44 | + | + | + | + | + | + | + | + | + | + | + | + |

### fMRI study

#### Behavioural responses

Some examples of typical verbal responses provided for each condition are shown in Table 2. Here one can clearly see the qualitative differences in response content across conditions. The autobiographical response is rich in spatio-temporal contextual details, typically including “what”, “where” and “when” information. In contrast the semantic-event condition elicits more generalised and schema-like event descriptions that are non-specific to a particular event. Finally, the semantic-object condition tends to include object-feature descriptions rather than events.

**Table 2.**
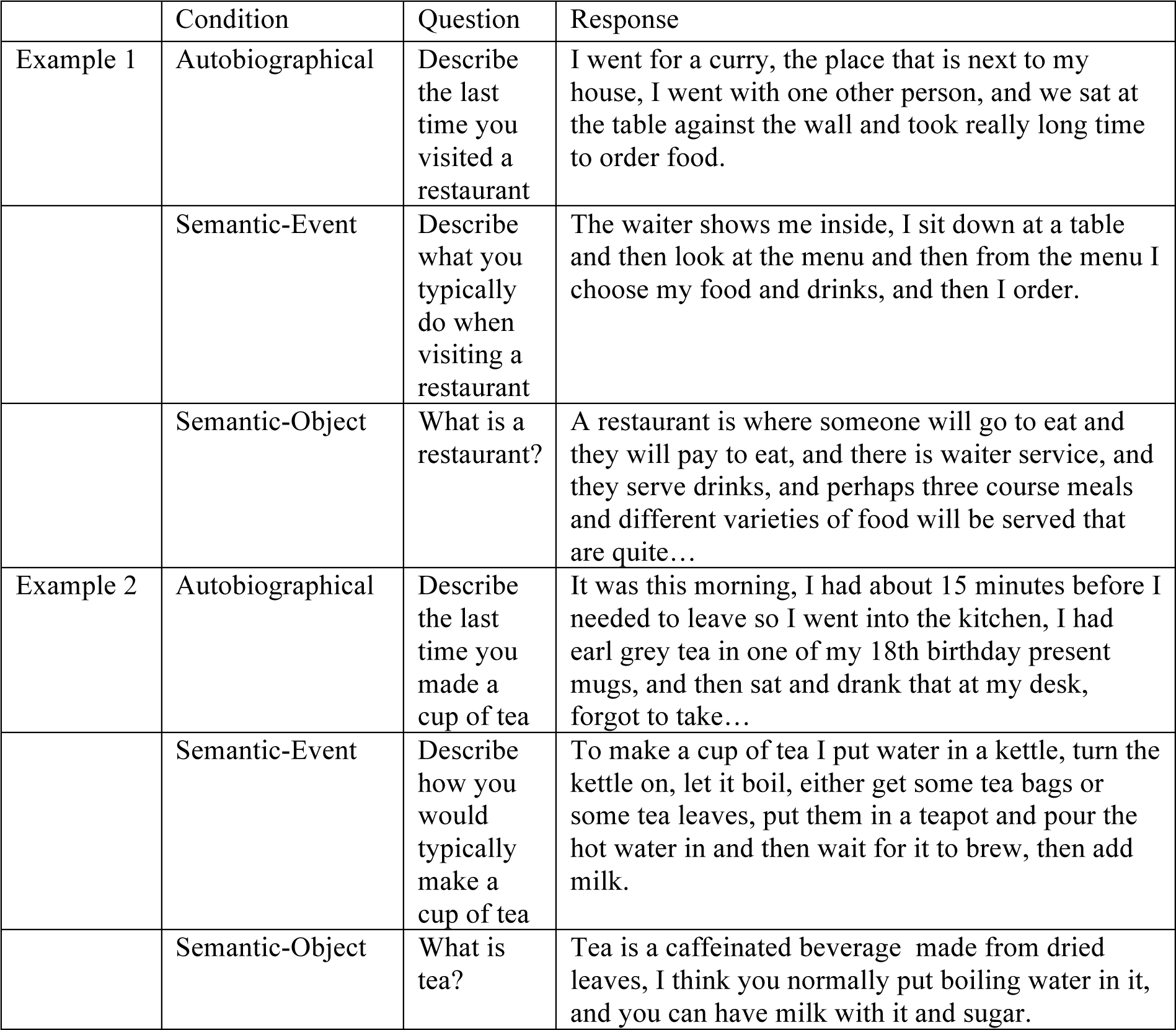
A typical example of a verbal response to each condition.

The participants were instructed to continue speaking for the entire 14 seconds. To check for difference in the quantity of speech content we compared the mean utterance length across conditions. The semantic-event condition elicited the longest responses (mean number of characters (SD) = 199.5 (34.78)) compared to the autobiographical condition (mean = 177.4 (35.33); *t*(37), = 7.24, *p* < .001), and the semantic-object condition (mean = 174.5 (35.80); *t*(37), = 10.79, *p* < .001). The autobiographical responses and semantic-object condition did not differ *(t*(37), = 0.93, *p* = .18). Participants produced the least amount of speech in the control condition compared to all experimental conditions (mean = 137.2 (35.33); all *t*s > 4.46, all *ps* < .001). Therefore, if results are to be explained purely in terms of quantity of speech produced one would expect the greatest activation for the semantic-event condition, whereas the autobiographical and semantic-object conditions would not differ.

#### fMRI results

##### Autobiographical retrieval vs. semantic retrieval

During the memory retrieval phase, all experimental conditions were found to activate a widespread left fronto-temporo-occipital network. In addition, the autobiographical condition was found to strongly activate the left AG (PGa and PGp), as well as other areas that are known to form the default mode network (medial fronto-parietal structures and hippocampus). The semantic-event condition did weakly activate the AG, however, the cluster was comparably smaller and limited to the anterior portion of AG, and there was little involvement of the wider DMN (Figure 2, Table S6). Figure S5 illustrates the locus of activation for both conditions relative to PGa and PGp, as defined by Caspers et al. (2008). Further analyses showed that this activation was largely driven by responses that contained greater autobiographical detail (see below). The semantic-object condition showed no pAG (or DMN) activation. Direct contrasts between conditions showed significantly stronger pAG and DMN activation for the autobiographical condition compared to either semantic task (Figure 2, Table S6). During the response phase, all tasks showed activation in areas associated with motor output (frontal lobe, insular, basal ganglia) as well as language areas in the temporal lobe. As per the results from the memory retrieval period, during the response phase the pAG was activated for the autobiographical task (and no other task), although the level of activation was lower and the cluster reduced in size compared to the memory retrieval phase (see Figure 2 showing whole brain results for each phase (see Figure S3 for comparable results comparing each experimental task > control task), and Figure 3 showing the ROI results for the left pAG, vATL, and autobiographical memory ROI to compare the signal magnitude of each phase). Indeed, an examination of the activation time-course showed that, whilst the pAG was activated throughout the question and response phase of the trial, it was more strongly associated with the memory retrieval phase, rather than response phase of the trial thereby suggesting a primary role in autobiographical memory retrieval. Accordingly, the remainder of the analyses focussed on the memory retrieval phase of the experiment (although comparable analyses from the response phase can be seen in Supplementary S7). It should be noted that whilst we opted to separate the retrieval and response phase of each trial, the overall pattern of results is the same when modelling the two phases together.

**Figure 2:**
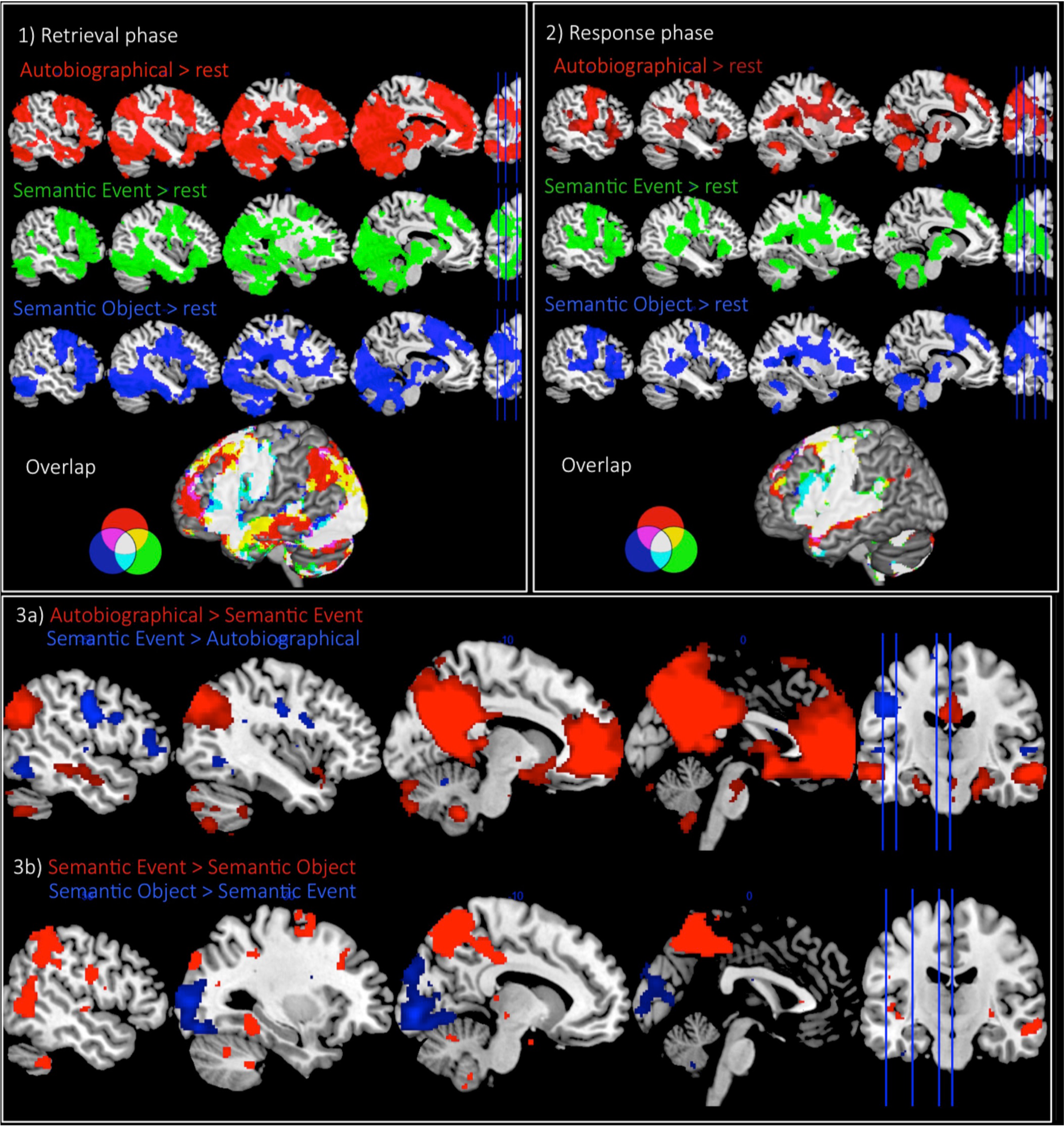
*Panel 1* - The fMRI results from each task > rest and the overlap in activation from the “retrieval phase”. *Panel 2* - The fMRI results from each task > rest and the overlap in activation from the “response phase”. *Panel 3* - a) The direct task contrasts of autobiographical condition > semantic event condition (red) (and vice versa in blue); b) The direct task contrasts of the semantic event condition > semantic object condition (red) (and vice versa in blue). Voxel height threshold p < .001, cluster corrected using FWE p < .05. MNI slice number *x* = -50, -40, -25, -10.

**Figure 3:**
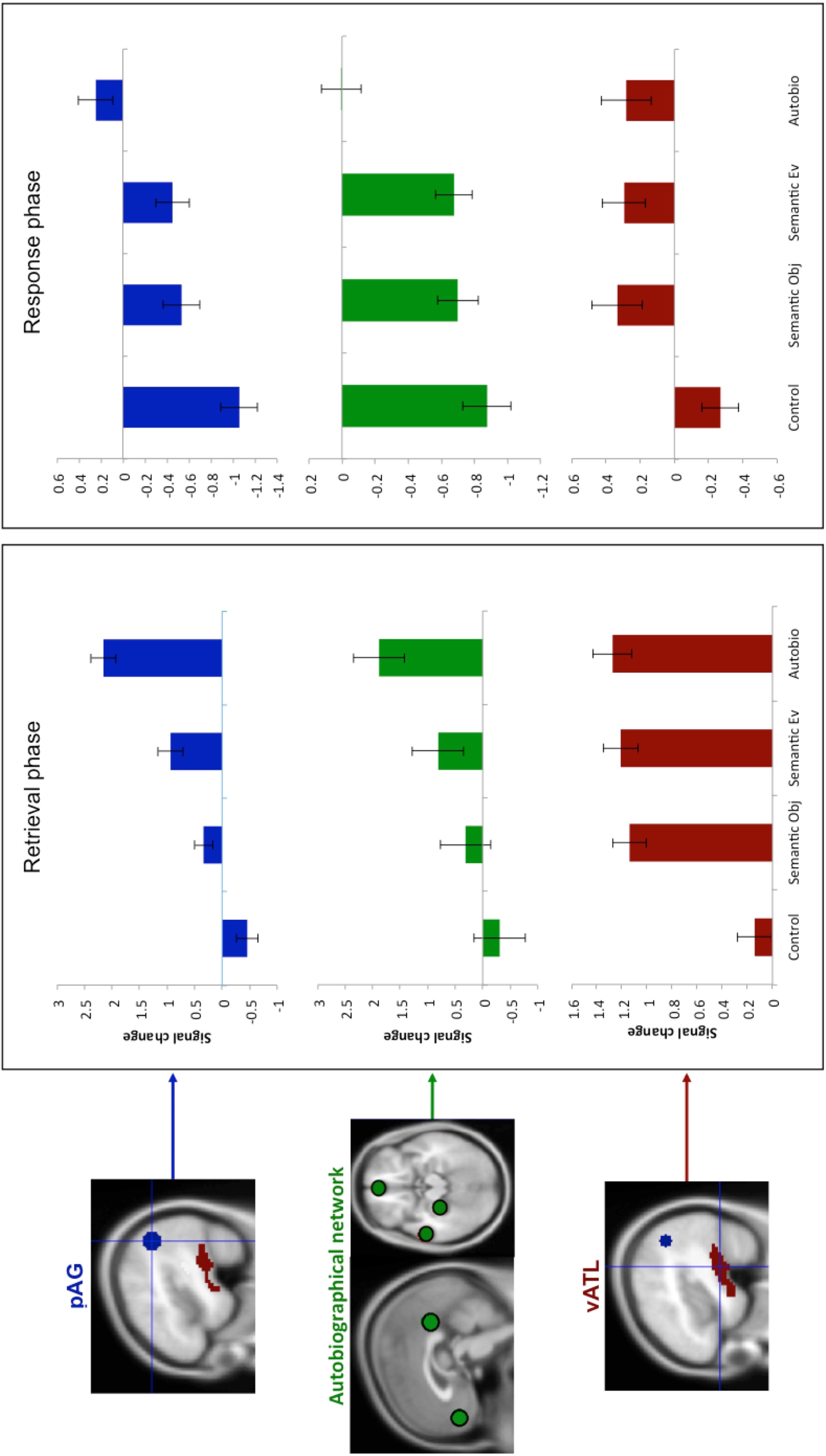
ROI results showing the signal change for each condition relative to rest in left pAG (blue), averaged across the wider left hemisphere autobiographical network (green), and left vATL (red).

##### Semantic-events vs. semantic objects

Directly contrasting the event-semantic and object-semantic tasks showed significantly stronger recruitment for semantic-event descriptions within the posterior middle temporal gyrus (pMTG), an area associated with multiple functions (motion-processing, verb processing, semantic control), and superior parietal lobe (SPL), whereas object-semantics showed increased activation of visual cortex, which could reflect the use of visual imagery during object description (Figure 2, Table S6).

#### ROI analyses

ROI analyses were conducted in order to compare, more closely, the response profile of the left pAG, vATL and the autobiographical memory network.

pAG: The pAG showed a strong positive activation for the autobiographical condition relative to rest (t(27) = 9.54, p < .001), and significantly greater activation relative to all other conditions (all ts > 8.00, all ps < .001). pAG also showed significant positive activation relative to rest for the semantic-events (t(27) = 4.13, p < .001), as well as greater activation for the semantic-events relative to semantic-objects. The semantic-objects did not differ significantly from rest (t(27) = 2.18, p = .06) but did from control (t(27) = 5.07, p < .001) since the control task deactivated the pAG (t(27) = -2.35, p = .02)) (all ts > 3.77, all ps < .001) (Figure 3).

vATL vs. pAG: the vATL showed a different response to the pAG across conditions, as illustrated by a significant Task x ROI interaction in a within-subject ANOVA (F(3) = 39.81, p < .001). Indeed, whereas the pAG responded differently to each experimental condition, the vATL showed equally strong involvement for all experimental conditions (all ts < 1.76, all ps > .09), which was significantly greater than rest (all ts > 8.22, all ps < .001), and control (all ts > 10.26, all ps < .001) (Figure 3). This is consistent with the role of the vATL as a semantic-hub, being activated by any task requiring semantic retrieval.

Autobiographical memory network vs. pAG: In contrast to the vATL, the autobiographical memory network showed a similar pattern of activation to that of the pAG. A Task x ROI within-subject ANOVA showing no significant interaction between regions (F(3) = 1.78, p > .19), and pairwise comparisons confirming that the autobiographical network responded in a similar pattern across conditions (all ts > 4.48, all ps < .001) (see Figure 3).

Together these results show that the pAG is not primarily involved in semantic memory retrieval similar to the vATL, but is more likely involved in some manner in autobiographical memory processes.

#### Abbreviations

Control (control non-propositional speech production task); Semantic Obj (semantic object definitions); Semantic Ev (Event definitions); Autobio (autobiographic descriptions)

#### Regression analyses

The GLM results demonstrate that the pAG, like other regions of the autobiographical network, was strongly activated by the autobiographical memory task. There was also some evidence of pAG activation for the event-semantic task, albeit at a lower magnitude. This is a very different pattern to that observed in the semantic memory network (including the vATL) where equal activation was found for all tasks. Whilst the pAG was clearly activated during autobiographical memory retrieval, what was driving the limited activation for the event-semantic task? The results of four regression analyses indicated that the pAG response to the event-semantic task was driven by the extent to which the participants were relying on an autobiographical retrieval strategy to respond to each question, rather than a semantic retrieval strategy. Specifically, we hypothesized that a novel and highly vivid event is less likely to be semanticised and more likely to rely on autobiographical memory retrieval. Consistent with this hypothesis, ROI analyses showed that pAG activation was: 1) positively related with event novelty (t(27) = 3.89, p = .001), and 2) positively with event vividness (t(27) = 3.81, p = .004). Furthermore, 3) there was a positive relationship with the “episodic memory strategy” variable (participants’ subjective rating of the extent they based their response on a specific memory rather than general knowledge; t(27) = 5.23, p < .001). Finally, 4) a negative relationship was found between pAG activation with the response similarity ratings, as computed by the LSA semantic similarity analysis (t(27) = -6.75, p = .001). This pattern can be clearly seen when conducting a median split analysis where high and low ratings are entered as separate variables in the GLM. This showed that as the item becomes more autobiographical in nature the activation increases, closer to that of the autobiographical task (Figure 4). Together these results suggest that the pAG is engaged when an event is based on autobiographical rather than semantic memory. Compellingly, showing a similar pattern as the AG, the wider autobiographical network was also significantly sensitive to each rating scale (all ts > 2.34, ps < .008), and the pAG and autobiographical network showed no ROI x rating scale interaction (F(3) = 2.22, p > .09) thereby indicating a similar response profile across regions. This result is in sharp contrast to that found in semantic areas, such as the vATL, which were invariant to each of these variables, showing no significant response regardless of the ratings (all ts < .6, ps > .6). Indeed, the response of the vATL was significantly different to the response of the pAG as shown by a significant region x rating scale interaction (F(3) = 46.17, p < .001). The fact that the pAG results mirror that of the episodic rather than semantic network provides strong evidence in favour of an episodic/autobiographical rather semantic function of the pAG in memory retrieval.

**Figure 4:**
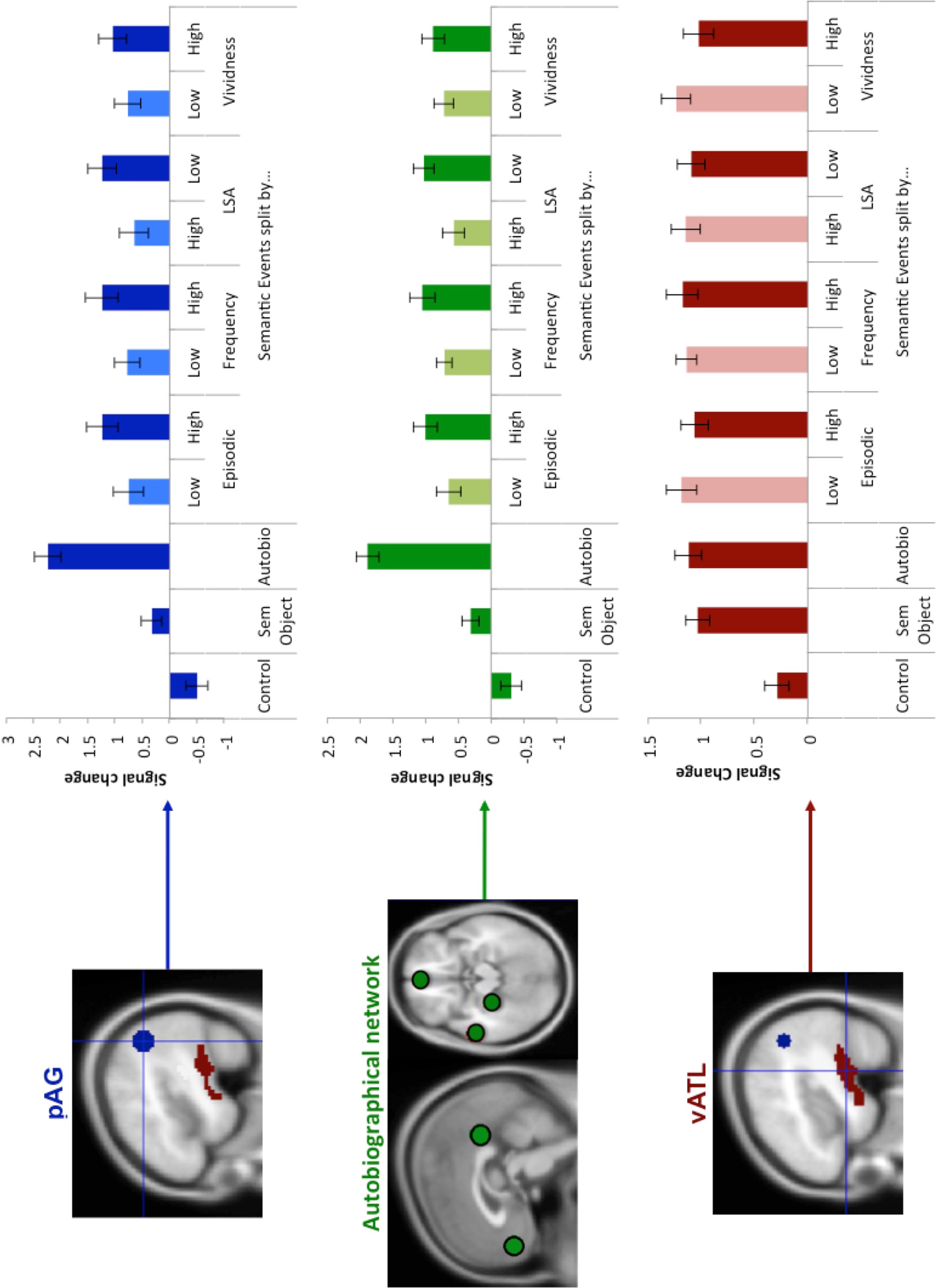
ROI results showing the signal change for each condition relative to rest in left pAG (blue), averaged across the wider left hemisphere autobiographical network (green), and left vATL (red). The Semantic-Event condition is split into a high vs. low condition based on a median-split using the ratings from Episodic relatedness, Frequency (novelty), LSA (response similarity), and Vividness analyses. *Abbreviations*: Control (control non-propositional speech production task); Sem Object (Object definitions)

## Discussion

The current study focused on comparing the (left) pAG engagement with three types of memory using the previously under-examined naturalistic task of propositional speech production. The results from the meta-analysis indicated that the pAG is only engaged as part of the propositional speech network when the task carries an autobiographical component. This conclusion was strongly supported by the results from the fMRI study. The key fMRI findings were: 1) pAG is positively engaged during autobiographical memory retrieval and activation increases with the degree to which a task relies on input from the episodic memory system. Elsewhere we have shown that the same pAG ROI, within area PGp, shows strong functional and structural connectivity with the episodic network (Humphreys et al., 2020; Humphreys et al., 2022) and indeed in the current study, the response profile across conditions in the pAG was found to be a very similar to the wider autobiographical network. 2) In contrast, the pAG showed a very different response profile to the vATL semantic hub – whilst pAG activation increased with the autobiographical nature of the task, the vATL was equally responsive to all conditions. This provides clear evidence that the pAG is not acting as a semantic hub. 3) Activation was strongest during the memory retrieval phase rather than the response phase of the trial.

### Implication for episodic research

The current data are consistent with a wealth of studies showing positive pAG (PGp) activation during autobiographic/episodic retrieval tasks, with activation correlated with memory vividness (Humphreys et al., 2022; Kuhl & Chun, 2014; Richter, Cooper, Bays, & Simons, 2016; Tibon, Fuhrmann, Levy, Simons, & Henson, 2019). We propose that information stored elsewhere in the episodic system is temporally buffered online in the pAG during episodic retrieval/memory construction. Indeed, the notion of an pAG episodic buffer has been proposed before (Vilberg & Rugg, 2008; A. D. Wagner et al., 2005), and fMRI evidence has shown that, unlike the hippocampus that responds to boundaries between episodic events, pAG activation is maintained throughout an event, consistent with the role of the pAG as an online buffer of retrieved information (Baldassano et al., 2017; Ben-Yakov & Henson, 2018; Van der Linden, Berkers, Morris, & Fernández, 2017). The buffer model is also consistent with neuropsychological evidence showing that, unlike medial temporal damage, parietal damage does not result in profound amnesia but rather it results in a lack of clarity or vividness of autobiographical details, which one might expect from a deficit in the buffering of multi-modal contextual information (Berryhill, Phuong, Picasso, Cabeza, & Olson, 2007; Bonnici, Cheke, Green, FitzGerald, & Simons, 2018; Davidson et al., 2008; Humphreys et al., 2021; Moscovitch, Cabeza, Winocur, & Nadel, 2016; Shimamura, 2011; Simons et al., 2008; St. Jacques, 2019; Yazar, Bergstrom, & Simons, 2014).

The current findings also speak to the notion that there is no straightforward dichotomy between episodic and semantic memory retrieval, rather the two exist along a continuum (Renoult, Irish, Moscovitch, & Rugg, 2019). This can be illustrated most clearly within the semantic-event condition, where there was a linear change in activation depending on the extent to which a memory pulled on the episodic vs. semantic systems - activation was modulated by event novelty, vividness, and level of semanticisation. This suggests that over time, as experiences reoccur, the memory for that event becomes less vivid and more generalizable and schema-like in nature, that reflect abstracted commonalties across experiences and are independent from the original spatio-temporal context (Gilboa & Marlatte, 2017).

### Implication for PUCC

The current data are consistent with the PUCC model, which assumes that (i) the entire lateral parietal cortex (LPC) may be underpinned by the same local computation which results in a domain general mechanism that can buffer temporally-extended information (as per ‘Elman’ PDP models) and that (ii) the buffered information varies across the LPC according to the proximity and connections to different inputs and outputs (Humphreys et al., 2020; Humphreys et al., 2022; Humphreys & Lambon Ralph, 2015; Humphreys et al., 2021). This tenet has been formally demonstrated in computational models, whereby the resultant “expressed behaviour” of a group of processing units depends not only on their local computation, but also on the long-range connectivity (more recently referred to as ‘connectivity-constrained cognition – C^3^’: (Chen, Lambon Ralph, & Rogers, 2017; Lambon Ralph, McClelland, Patterson, Galton, & Hodges, 2001; Plaut, 2002)). Indeed, we have proposed two primary axes of LPC organisation: dorsal-ventral and anterior-posterior. Driven by top-down input from the prefrontal cortex (Crowe et al., 2013), the dorsal LPC is actively engaged by executively demanding tasks across cognitive domains (Fedorenko, Duncan, & Kanwisher, 2013; Humphreys & Lambon Ralph, 2015, 2017a) and may be involved in the selection and/or manipulation of buffered information, akin to an executively-penetrated working memory system (Humphreys & Lambon Ralph, 2015; Pessoa, Gutierrez, Bandettini, & Ungerleider, 2002). In contrast, we have shown that in the ventral LPC there is an anterior-posterior gradient of activation, with more anterior regions (in PGa) positively engaged during sentence/narrative comprehension and showing structural/functional connectivity with posterior temporal language-related areas, pAG (within anterior PGp) is positively engaged during episodic/autobiographical and this is driven by strong functional and structural connectivity with the episodic memory system, and more posterior LPC (posterior PGp) showing positive engagement for picture-sequences, and connecting with visual cortex (Humphreys et al., 2020; Humphreys et al., 2022). When a task does not involve the cognitive process of interest, that particular ventral LPC subregion is deactivated – hence the common observation of AG task-related deactivation.

A system concerned with temporal buffering would be particularly useful for tasks that are reliant on information that unfolds over time. This includes autobiographical retrieval, which is considered to require active re-construction and “re-living” (Conway et al., 1999; Tulving, 2002). Indeed, parietal damage can produce deficits in performing time-extended verbal and non-verbal behaviours (e.g., logopenic progressive aphasia, ideational apraxia, etc.) (Buxbaum, Kyle, Tang, & Detre, 2006; Gorno-Tempini et al., 2008). Furthermore, the anterior AG (PGa) is engaged by sentence processing and narrative speech comprehension (Branzi, Humphreys, Hoffman, & Lambon Ralph, 2020; Branzi, Pobric, Jung, & Lambon Ralph, 2021; Humphreys et al., 2020; Humphreys & Lambon Ralph, 2015) and a more posterior area of AG (posterior PGp) shows a reliable response to narratives and movies when the content is intact rather than temporally scrambled (Hasson, Yang, Vallines, Heeger, & Rubin, 2008; Lerner, Honey, Silbert, & Hasson, 2011). The indication that the AG is more strongly activated during the memory-retrieval phase than the response phase of the trial has implications for the nature of the PUCC buffering system. Specifically, these findings are more consistent with the AG acting as an “input” rather than “output” buffer whereby LPC buffers temporally-extended external and internal inputs, rather than controlling or monitoring the roll out of temporally-extended behaviours. Indeed, the finding that the system was not activated during speech output is consistent with evidence that the area is not reliably activated by other types of speech output tasks, e.g., verbal fluency (S. Wagner, Sebastian, Lieb, Tüscher, & Tadić, 2014). Furthermore, outside of speech production, the AG has been found to be more active when imagining the words to a familiar song than listening to the same song, supporting the notion that the region is sensitive to the processes involved in memory retrieval, rather than the external stimulus itself (Herholz, Halpern, & Zatorre, 2012).

In terms of a mechanistic account of how the buffer might operate, two possible alternatives can be contemplated. First, temporally-extended behaviours are often thought to rely upon a stored “task-schema” (Bornkessel-Schlesewsky & Schlesewsky, 2013; Rumelhart, Smolensky, McClelland, & Hinton, 1986; Schmidt, 1975). Accordingly, the buffer might keep track of not only the evolving state of play but also the remaining steps needed to transform a particular “thought” into speech output (the future steps). Given that the AG appears to be differentially activated for the retrieval vs. speech phase of autobiographical production, then this schema-based hypothesis seems less likely. Alternatively and consistent with the current data, the AG might only buffer incoming spatio-temporally varying information, and does not contain information with regard the future steps required to transform the “thought” into action (Botvinick & Plaut, 2004, 2006). This approach allows for greater flexibility in real-world unpredictable situations, since an ever-changing context will lead to variability in the outcome of a sequence each time it is performed.

### Implications for semantic research

The current data provide clear evidence that the pAG is not engaged during semantic retrieval, supporting findings elsewhere (Humphreys et al., 2022; Humphreys & Lambon Ralph, 2015, 2017a). This is consistent with evidence showing that parietal damage does not result in gross semantic-memory retrieval impairment (unlike damage to the vATL), but rather in the flexible use and manipulation of semantic information (Jefferies & Lambon Ralph, 2006). The current data also go one-step further by demonstrating that the pAG is not sensitive to all semantic-events, rather activation is driven by the extent by which event-retrieval relies on input from the autobiographical/episodic memory system. It is important to note, however, that beyond PGp, other areas of LPC may be sensitive to semantic information in cases that require the buffering of temporally-extended information. For instance, anterior-ventral IPL (bordering with temporoparietal junction) responds to sentence processing and narrative speech comprehension (Branzi et al., 2020; Branzi et al., 2021; Humphreys et al., 2020; Humphreys & Lambon Ralph, 2015), and more ventral and posterior regions respond to picture sequences or semantic-association tasks using pictures (Humphreys et al., 2020; Seghier, Fagan, & Price, 2010).

How can one align the lack of semantic modulation, found here and elsewhere, with neuroimaging results showing an apparent sensitivity to semantic contrasts or semantic-events in the pAG (Binder & Desai, 2011; Binder et al., 2009)? As mentioned in the Introduction, when studying LPC function it is important to take into account certain critical variables: 1) the polarity of activation relative to rest and 2) difficulty-related differences. With regard the polarity of activation, direct comparisons have shown that, whereas the pAG is positively engaged during episodic retrieval, the same region is deactivated relative to rest during semantic retrieval tasks (Humphreys et al., 2022; Humphreys & Lambon Ralph, 2015, 2017a). It is of course difficult to interpret “rest”, as it may involve spontaneous semantic and linguistic processing (Binder et al., 1999), although the same argument would presumably also apply to episodic retrieval which shows positive task-related activation (Andrews-Hanna, 2012; Buckner et al., 2008). Critically, however, the pAG semantic-hub hypothesis is inconsistent with the observation that the pAG shows a very different pattern to other semantic-hubs, such as the vATL, which is *positively* engaged during semantic retrieval (Humphreys et al., 2015; Humphreys & Lambon Ralph, 2017a), and shows constant activation across conditions in the current study (presumably due to the fact that all conditions contain meaningful elements and entities, and speech production is driven by semantics). Nevertheless, of course, it is possible that in some circumstances, there may be certain aspects of a semantic-retrieval task that do modulate AG, but critically not in the way currently specified by existing semantic models. With regard to task-difficulty, the contrast of word > nonword and concrete > abstract typically involve comparing an easier task vs. a harder task, and pAG deactivation is known to relate to task difficulty, with increasing deactivation for harder tasks (Gilbert, Bird, Frith, & Burgess, 2012; Hahn, Ross, & Stein, 2007; Harrison et al., 2011; Humphreys & Lambon Ralph, 2017a). Indeed, direct examinations of task difficulty in semantic and non-semantic tasks have revealed a main effect of task difficulty (easy vs. hard) on the degree of deactivation in the pAG but no semantic vs. non-semantic difference (Humphreys & Lambon Ralph, 2017b; Quillen, Yen, & Wilson, 2021), and compellingly, the classic pattern of differential activation associated with the contrast of words > non-words or concrete > abstract processing, can be flipped by reversing the difficulty of the task or the stimuli (Graves, Boukrina, Mattheiss, Alexander, & Baillet, 2017; Pexman, Hargreaves, Edwards, Henry, & Goodyear, 2007). A similar argument might also explain results from studies of combinatorial-semantics, where ‘higher’ pAG ‘activation’ is associated with adjective-noun or adjective-verb word pairs that are readily-combined compared to less automatic combinations (Bemis & Pylkkanen, 2013; Matchin, Liao, Gaston, & Lau, 2019).

### Implications for models of speech production

The current meta-analysis constitutes the largest meta-analysis of the propositional speech production literature. Together with the results from the fMRI study, the data have implication for models of speech production. Specifically, the results show that the core speech production network is made up of frontal areas (IFG, precentral gyrus, and SMA), and temporal areas (middle temporal gyrus, and the vATL). In contrast, the pAG plays a task-specific role. Several researchers have proposed a dual-stream model of language, where the neural mechanisms that underlie language can be separated into dorsal and ventral processing streams (Hickok & Poeppel, 2007; Ueno, Saito, Rogers, & Lambon Ralph, 2011; Weiller, Bormann, Saur, Musso, & Rijntjes, 2011). Perhaps the best-known of the dual-stream models proposes that language production and comprehension rely on separable dorsal and ventral processing streams, respectively (Hickok & Poeppel, 2007). Here, the ventral route consists of posterior and ventral temporal cortex for receptive speech, whereas the dorsal stream consists of dorsal posterior temporal cortex and posterior frontal areas, with the frontal cortex serving a primarily a motoric role in speech output. The current data show that both dorsal and ventral fronto-temporal networks are implicated in the core speech production system. More specifically, we found extensive frontal activation during speech production. However, we argue for a broader role of frontal cortex in speech production beyond the obvious motor component of the task, such as executive control and working memory processes. Indeed, lateral frontal areas extending into ventral IFG are known to be involved in semantic control of speech comprehension and production as well as nonverbal tasks (Hoffman, 2019; Hoffman, Jefferies, & Lambon Ralph, 2010; Jackson, 2021; Lambon Ralph, Jefferies, Patterson, & Rogers, 2017), and areas overlapping with the multi-demand network (DLPFC and SMA) are involved in domain-general executive processing (Fedorenko et al., 2013). In terms of the contribution of the temporal lobe, the full length of the MTG and ventral vATL were strongly activated across conditions in the fMRI study. The posterior MTG has been associated with semantic control (Jackson, 2021; Lambon Ralph et al., 2017). In contrast, the more anterior MTG is known to be active during sentence-level tasks during speech production and comprehension (Branzi et al., 2020; Hoffman, Binney, & Lambon Ralph, 2015; Humphreys & Gennari, 2014; Morales, Patel, Tamm, Pickering, & Hoffman, 2022), and may play a role in combinatorial processes (Humphries, Binder, Medler, & Liebenthal, 2006). In terms of the vATL, there is strong evidence that the vATL is multi-modal semantic hub (Lambon Ralph et al., 2017). For instance, the contribution of the vATL to speech production and comprehension is well-established in the neuropsychological literature: semantic dementia patients show significant deficits in both expressive and receptive semantic tasks (Jefferies & Lambon Ralph, 2006) including impoverished content in propositional speech tasks (Bird, Lambon Ralph, Patterson, & Hodges, 2000; Hoffman, Meteyard, & Patterson, 2014) Furthermore, vATL engagement during semantic tasks has been consistently shown using fMRI and TMS in healthy participants and intracortical grid electrode studies in neurosurgical patients (Binney, Embleton, Jefferies, Parker, & Lambon Ralph, 2010; Humphreys & Lambon Ralph, 2015; Rogers et al., 2021; Shimotake et al., 2014; Visser, Jefferies, & Lambon Ralph, 2010). Reliable vATL activation was less obvious in the meta-analysis (although still present at a reduced threshold). This absence of evidence in the meta-analysis is very likely to be due to various methodological issues including fMRI signal loss in most single-echo EPI fMRI paradigms (Devlin et al., 2000; Visser et al., 2010). The current fMRI study utilised a multi-echo paradigm designed to improve signal coverage thereby revealing strong vATL modulation across semantic conditions (Halai, Welbourne, Embleton, & Parkes, 2014)

## Conclusions

To conclude, we demonstrated here that left pAG activation depends on the extent to which a task requires input from the autobiographical-, rather than semantic-memory retrieval systems. The results are consistent with a model in which the LPC acts as a multi-modal buffer and subregional variations in expressed functions reflect differences in their long-range connectivity; the left pAG gravitates to episodic-related tasks as a result of its preferential connectivity to the episodic network.

## Acknowledgements

This research was supported by an MRC Programme grant to MALR (MR/R023883/1) and an intramural award (MC_UU_00005/18). The authors declare no conflict of interest.

## Open access

For the purpose of open access, the UKRI-funded authors have applied a Creative Commons Attribution (CC BY) licence to any Author Accepted Manuscript version arising from this submission.

## Data availability statement

The data will be made available at http://www.mrc438cbu.cam.ac.uk/publications/opendata

## Author contributions

G. F. H. and M. A. L. R. conceived and planned the experiments. G. F. H. performed the meta-analysis and collected the fMRI data. G. F. H. and A. D. H. performed the fMRI data analysis. G. F. H. and F. M. B. coded the speech recordings and performed the latent semantic analysis. G. F. H. and M. A. L. R. interpreted the results and wrote and revised the manuscript. All authors provided critical feedback on the manuscript.

