## Supplementary for "The left posterior angular gyrus is engaged by autobiographical recall not object-semantics, or event-semantics: Evidence from contrastive propositional speech production"

Table S1. A description of the studies included in the metaanalysis.

*An "unconstrained" task is that in which the participants were free to respond in any way they choose, in contrast to a "constrained" tasks whereby participants were restricted in how they responded (e.g. the response must avoid certain words).

| Study | Task | Contrast | Technique | N. Participants | Activated the AG |
|---|---|---|---|---|---|
| Braune et al. 2001 | Generate a personal story | Speech > oral-motor baseline | PET | 12 | Yes |
| Blank, Scott, Murphy, Warburton, and Wise (2002) | Generate a personal story | Speech > counting + nursery rhyme recitation | PET | 8 | Yes |
| Haller, Radue, Erb, Grodd, and Kircher (2005) | Generate a sentence based on three words | Sentence generation > word reading | fMRI | 15 | No |
| | | Sentence generation > sentence reading | | | No |
| Kemeny, Ye, Birn, and Braun (2005) | Generate a sentence based on verb | Sentence generation > syllable repetition (pa-ta-ka) | fMRI | 6 | No |
| Troiani et al. (2008) | Generate a story based on pictures | Sentence generation > story viewing + syllable repetition (pa-da-ka) | fMRI | 15 | No |
| Tremblay and Small (2011) | Object description | Sentence generation > picture viewing | fMRI | 21 | No |
| Menenti et al. (2011) | Picture description | Repeated words in production > repeated words in comprehension | fMRI | 20 | No |
| Geranmayeh et al. (2012) | Unconstrained | Define nouns > tongue movement | fMRI | 19 | No |
| Grande et al. (2012) | Constrained (avoid frequent words) | Sentence generation > rest | fMRI | 18 | No |
| Menenti, Segaert, and Hagoort (2012) | Picture description | Novel semantics > repeated semantics | fMRI | 20 | No |
| | | Novel words > repeated words | | | |
| | | Novel syntax > repeated syntax | | | |
| Menenti, Petersson, and Hagoort (2012) | Picture description | Novel words > repeated words | fMRI | 24 | No |
| | | Novel meaning > repeated meaning | | | |
| Geranmayeh, Wise, Mehta, and Leech (2014) | Define a noun | Define a noun > counting | fMRI | 24 | No |
| Schönberger et al. (2014) | Generate a sentence | Generation of simple complete sentences > sentences missing verb | fMRI | 15 | Yes |
| Simmonds, Leech, Collins, Redjep, and Wise (2014) | Unconstrained | Overt noun definition > rest (look at a row of Xs) | fMRI | 17 | No |
| Matchin and Hickok (2016) | Active or passive | Sentence generation > word lists | fMRI | 20 | No |

| | | | | | |
|---|---|---|---|---|---|
| Dhanjal, Handunnetthi, Patel, and Wise (2008) | Define a noun | Define a noun > counting | fM RI | 21 | No |
| Brownsett and Wise (2010) | Self-referential narrative | Self-referential speech > syllables | PET | 13 | Yes |
| Humphreys and Gennari (2014) | Sentence completion | Sentence completion > reading | fM RI | 17 | No |
| Braun et al. (1997) | Self-referential speech | Production tasks > mouth movements | PET | 8 | Yes |
| Kircher et al. (2004) | Picture description | Free speech > rest | fM RI | 6 | No |
| Haller, Radue, Erb, Grodd, and Kircher 2005 | Sentence generation based on key words | Sentence generation > word reading | fM RI | 15 | No |
| Tremblay and Small (2011) | Generate sentence from pictures | Sentence generation > rest | fM RI | 21 | No |
| Kemeny, Ye, Birn, and Braun (2005) | Sentence generation based on verb | Sentence generation > rest | fM RI | 6 | No |
| AbdulSabur, Xu, Chow, Baxter, Carson, and Braun (2014) | Retell a previously presented story | Narrative production > nursery rhyme | fM RI/ PET | 18 | No |
| Yuan, Major-Girardin, and Brown (2018) | Free narrative production | Narrative production > picture description | fM RI | 24 | Yes |

Table S2. A complete list of experimental stimuli in include in the fMRI task.

| A list of experimental stimuli | | |
|---|---|---|
| Condition | ItemN | Question |
| autobio | 1 | Describe the last time you arranged a holiday |
| autobio | 2 | Describe the last time you used a bike |
| autobio | 3 | Describe your most recent journey to work/university |
| autobio | 4 | Describe the last time you bought a present |
| autobio | 5 | Describe the last time you caught a train |
| autobio | 6 | Describe the last time you sent an email |
| autobio | 7 | Describe the last time you brushed your teeth |
| autobio | 8 | Describe the last time you bought clothes or shoes |
| autobio | 9 | Describe the last time you made the bed |
| autobio | 10 | Describe the last time you took a photograph |
| autobio | 11 | Describe the last time you cleaned the dishes |
| autobio | 12 | Describe the last time you went to a shopping centre |
| autobio | 13 | Describe the last time you went to a restaurant |
| autobio | 14 | Describe the last time you went to the pub |
| autobio | 15 | Describe the last time you met with a friend |
| autobio | 16 | Describe the last time you sent a message using your phone |
| autobio | 17 | Describe the last time you had lunch |
| autobio | 18 | Describe the last time you caught a bus |
| autobio | 19 | Describe what you did for dinner last night |
| autobio | 20 | Describe the last time you went to a supermarket or food shop |
| autobio | 21 | Describe your most recent trip to a café |
| autobio | 22 | Describe the last time you went to a party |
| autobio | 23 | Describe the last time you used a kettle |
| autobio | 24 | Describe the last time you ate desert |
| autobio | 25 | Describe the last time you spoke on the phone |
| autobio | 26 | Describe the last time you went to the airport |
| autobio | 27 | Describe the last time you watched a TV show or movie |
| autobio | 28 | Describe the last time you had breakfast |
| autobio | 29 | Describe the last time you had coffee or tea |
| autobio | 30 | Describe the last time you had your haircut |
| autobio | 31 | Describe the last time you wrote an essay or paper |
| autobio | 32 | Describe the last time you cleaned the kitchen |
| autobio | 33 | Describe the last time you did the laundry |
| autobio | 34 | Describe the last time you used the oven |

| | | |
|---|---|---|
| autobio | 35 | Describe the last time you ate fruit or vegetables |
| autobio | 36 | Describe the last time you travelled by car |
| semEvent | 1 | Describe what you would typically do when visiting a restaurant |
| semEvent | 2 | Describe what you would typically do when visiting a pub |
| semEvent | 3 | Describe what you would typically do to arrange to meet with a friend |
| semEvent | 4 | Describe how you would typically write a text message |
| semEvent | 5 | Describe what you would typically do to make a jam sandwich |
| semEvent | 6 | Describe what you would typically do to catch a bus |
| semEvent | 7 | Describe what you would typically do to cook pasta |
| semEvent | 8 | Describe what you would typically do when visiting the supermarket |
| semEvent | 9 | Describe what you would typically do in a café |
| semEvent | 10 | Describe what you would typically do at a party |
| semEvent | 11 | Describe how you would typically use a kettle |
| semEvent | 12 | Describe how you would typically make a cake |
| semEvent | 13 | Describe how you would typically make a phone call |
| semEvent | 14 | Describe what typically happens when you visit an airport |
| semEvent | 15 | Describe what you would typically do when visiting the cinema |
| semEvent | 16 | Describe what you would typically do to fry an egg |
| semEvent | 17 | Describe how you would typically make a cup of tea |
| semEvent | 18 | Describe what typically happens when you go for a haircut |
| semEvent | 19 | Describe how you would typically write an essay |
| semEvent | 20 | Describe what you would typically do to clean the kitchen |
| semEvent | 21 | Describe how you would typically do the laundry |
| semEvent | 22 | Describe how you would typically use an oven |
| semEvent | 23 | Describe how you would typically chop vegetables |
| semEvent | 24 | Describe what you typically do to start driving a car |
| semEvent | 25 | Describe what you would typically do in order to arrange a holiday |
| semEvent | 26 | Describe how you would typically ride a bike |
| semEvent | 27 | Describe how you typically travel to work |
| semEvent | 28 | Describe what you would typically do in order to buy a present |
| semEvent | 29 | Describe what you would typically do when catching the train |
| semEvent | 30 | Describe what you would typically do to send an |

| | | |
|---|---|---|
| | | email |
| semEvent | 31 | Describe what you would typically do to brush your teeth |
| semEvent | 32 | Describe what you would typically do when clothes shopping |
| semEvent | 33 | Describe how you would typically make the bed |
| semEvent | 34 | Describe what you would typically do to take a photograph |
| semEvent | 35 | Describe how you typically cleans the dishes |
| semEvent | 36 | Describe what you would typically do at a shopping centre |
| semObj | 1 | What is a phone? |
| semObj | 2 | What is an airport? |
| semObj | 3 | What is a movie? |
| semObj | 4 | What is breakfast? |
| semObj | 5 | What is coffee? |
| semObj | 6 | What is a haircut? |
| semObj | 7 | What is an essay? |
| semObj | 8 | What is a kitchen? |
| semObj | 9 | What is laundry? |
| semObj | 10 | What is an oven? |
| semObj | 11 | What is fruit? |
| semObj | 12 | What is a car? |
| semObj | 13 | What is a holiday? |
| semObj | 14 | What is a bike? |
| semObj | 15 | What is a University? |
| semObj | 16 | What is a present? |
| semObj | 17 | What is a train? |
| semObj | 18 | What is an email? |
| semObj | 19 | What is a toothbrush |
| semObj | 20 | What are clothes? |
| semObj | 21 | What is a bed? |
| semObj | 22 | What is a photograph? |
| semObj | 23 | What is a dish? |
| semObj | 24 | What is a shopping centre? |
| semObj | 25 | What is a restaurant? |
| semObj | 26 | What is a pub? |
| semObj | 27 | What is a friend? |
| semObj | 28 | What is a text message? |
| semObj | 29 | What is lunch? |
| semObj | 30 | What is a bus? |

| | | |
|---|---|---|
| semObj | 31 | What is dinner? |
| semObj | 32 | What is a supermarket? |
| semObj | 33 | What is a café? |
| semObj | 34 | What is a party? |
| semObj | 35 | What is a kettle? |
| semObj | 36 | What is desert? |

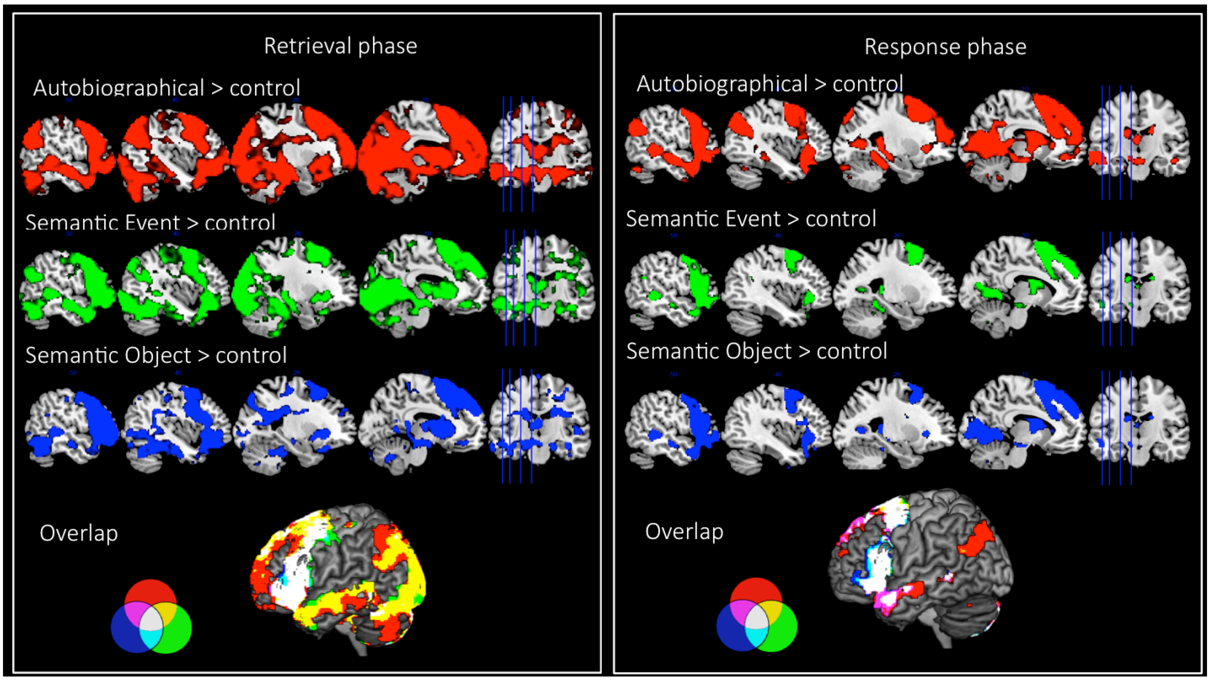


Figure S3. The fMRI results from each experimental task > control task from the "retrieval phase" (left), "response phase" (right) (yellow, white, cyan, violet reflect overlapping activations). Voxel height threshold p < .001, cluster corrected using FWE p < .05

Table S4. The peak coordinates from significant clusters in the metaanalysis of 25 speech production studies.

| Region | | Coordinates | | |
|---|---|---|---|---|
| Parietal cluster | SPL | -21 | -67 | 49 |
| | Posterior cingulate/precuneus | 2 | -57 | 12 |
| | IPS | -30 | -53 | 44 |
| | Precuneus | -7 | -59 | 29 |
| | Precuneus | -7 | -59 | 50 |
| | AG | -52 | -59 | 26 |
| | AG | -44 | -74 | 30 |
| Temporal clusters | MTG | -59 | -46 | 4 |
| | SMA | -4 | 12 | 59 |
| Frontal clusters | Precentral gyrus | -44 | 3 | 50 |
| | IFG BA44 | -47 | 20 | 8 |

Figure S5. The locus of activation for the contrast of Autobiographical > rest (red) and the Semantic event > rest (green), overlap (yellow) during the "retrieval phase" (as shown in Figure 2) relative to the anatomical boundaries of PGa (anterior) and PGp (posterior), as defined by Caspers et al. (2008).

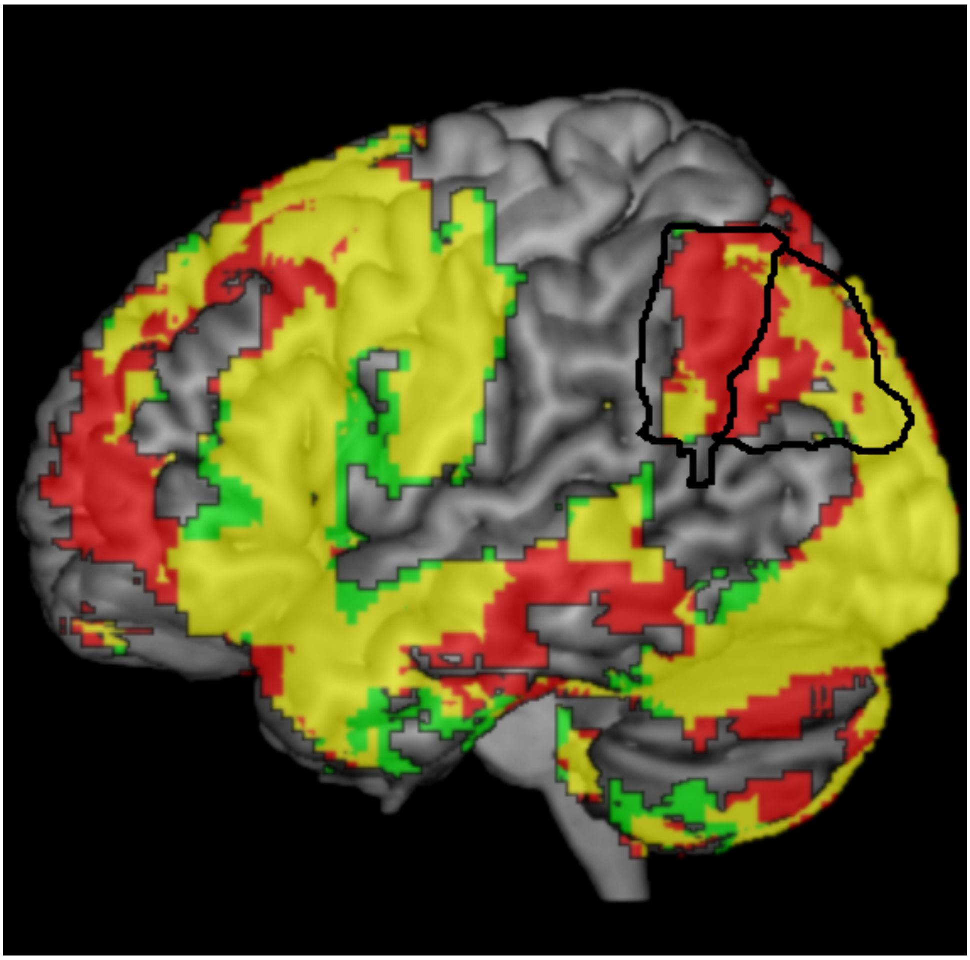

Table S6. The fMRI results from each task > rest and the overlap in activation from the “retrieval phase”. Voxel height threshold p < .001, cluster corrected using FWE p < .05

| Contrast | Cluster size | T | MNI coordinates |
|---|---|---|---|
| autobiographical > rest | 15988 | 18.83 | -16 -60 10 |
| | | 17.38 | 4 -72 -28 |
| | | 16.01 | 14 -84 -4 |
| | 1342 | 12.51 | -16 16 54 |
| | | 12.4 | -10 22 56 |
| | | 11.7 | -10 44 36 |
| | 267 | 12.45 | -30 34 -2 |
| | | 9.77 | -38 36 -10 |
| | | 9.18 | -36 46 0 |
| | 347 | 11.76 | -34 -68 38 |
| | | 10.03 | -46 -70 32 |
| | | 9.45 | -30 -78 42 |
| | 312 | 11.64 | -52 -2 24 |
| | | 10.36 | -46 -8 26 |
| | | 10.12 | -52 -10 36 |
| | 206 | 11.38 | -38 -50 20 |
| | | 11.27 | -44 -50 26 |
| | | 9.52 | -36 -48 28 |
| | 691 | 11.08 | 16 20 30 |
| | | 10.24 | 22 8 38 |
| | | 10.21 | 28 0 38 |
| | 121 | 11.05 | 32 44 -2 |
| | | 10.81 | 36 38 -6 |
| | 302 | 10.98 | 36 -30 -10 |
| | | 10.86 | 38 -18 -18 |
| | | 10.27 | 32 -12 -18 |
| | 189 | 10.4 | -50 18 18 |

| | | | | | |
|---|---|---|---|---|---|
| | | 9.16 | -42 | 16 | 40 |
| | | 9.04 | -40 | 14 | 24 |
| | 144 | 9.94 | -14 | -38 | 30 |
| | | 9.72 | -4 | -40 | 34 |
| | | 9.7 | -16 | -44 | 36 |
| object semantics > rest | 40374 | 14.13 | -12 | -62 | -22 |
| | | 13.7 | 34 | 36 | -4 |
| | | 13.56 | -6 | 10 | 58 |
| | 104 | 8.66 | 20 | -12 | -4 |
| | | 8 | 18 | -6 | 2 |
| | 125 | 8.08 | -10 | 44 | 34 |
| | | 7.79 | -8 | 52 | 32 |
| | | 6.87 | -8 | 42 | 46 |
| | 111 | 7.98 | -56 | -38 | 4 |
| | | 6.83 | -64 | -40 | 6 |
| event semantics > rest | 45385 | 15.73 | 6 | -74 | -26 |
| | | 15.19 | 34 | 36 | -4 |
| | | 14.33 | 14 | -86 | -4 |
| | 147 | 8.37 | -52 | -38 | 4 |
| | | 6.64 | -66 | -38 | 2 |
| | 33 | 7.93 | -22 | -20 | 42 |
| | | 6.43 | -18 | -20 | 50 |
| | 116 | 7.85 | 40 | 18 | -26 |
| | | 6.93 | 34 | 14 | -32 |
| | | 6.92 | 32 | 12 | -40 |
| Autobiographical > event semantics | 11415 | 11.2 | 0 | -64 | 28 |
| | | 10.47 | -10 | -58 | 30 |
| | | 10.13 | 4 | -56 | 28 |
| | 11875 | 10.02 | -4 | 58 | -6 |
| | | 9.09 | -6 | 40 | -4 |
| | | 8.53 | -4 | 54 | 16 |

| | | | |
|---|---|---|---|
| | 1332 | 6.71 | -46 -60 30 |
| | | 6.22 | -44 -74 42 |
| | | 5.37 | -52 -70 30 |
| | 991 | 6.33 | -58 -14 -14 |
| | | 5.66 | -58 -6 -18 |
| | | 5.47 | -68 -16 -14 |
| | 1529 | 6.28 | 60 -8 -16 |
| | | 5.98 | 52 16 -32 |
| | | 5.86 | 30 20 -16 |
| | 1344 | 6.08 | 50 -60 26 |
| | | 5.59 | 44 -72 40 |
| | | 5.25 | 52 -72 22 |
| | 550 | 5.78 | -4 -52 -46 |
| | | 5.76 | 10 -48 -44 |
| | | 4.72 | 0 -58 -50 |
| | 261 | 5.37 | -30 10 -16 |
| | | 4.28 | -38 14 -26 |
| | | 3.26 | -40 14 -12 |
| | 580 | 4.63 | -28 -84 -36 |
| | | 4.45 | -40 -78 -38 |
| | | 4.19 | -8 -84 -38 |
| | 274 | 4.55 | 26 -86 -28 |
| | | 3.99 | 6 -86 -20 |
| | | 3.96 | 38 -86 -30 |
| Semantic event > autobiogrphical | 1266 | 5.53 | -48 -14 38 |
| | | 5.46 | -52 -8 28 |
| | | 5.09 | -58 -8 18 |
| | 970 | 5.12 | 62 -6 24 |
| | | 5.07 | 52 -2 24 |
| | | 4.94 | 52 -8 30 |
| | 178 | 4.39 | -48 34 2 |

| | | | |
|---|---|---|---|
| | 265 | 4.13 | -50 -62 -12 |
| | | 3.57 | -42 -54 -6 |
| | | 3.45 | -42 -50 -14 |
| Semantic event > semantic Object | 4240 | 8.32 | 8 -60 54 |
| | | 8.27 | -8 -68 50 |
| | | 8.24 | -10 -62 58 |
| | 2739 | 8.15 | 26 14 58 |
| | | 6.52 | 32 6 66 |
| | | 5.47 | 36 26 52 |
| | 3024 | 7.59 | 56 -40 46 |
| | | 6.87 | 56 -50 20 |
| | | 6.67 | 60 -48 32 |
| | 1328 | 6.83 | -60 -38 44 |
| | | 4.07 | -48 -46 60 |
| | | 3.97 | -34 -42 42 |
| | 801 | 6.78 | -24 4 60 |
| | | 4.77 | -24 -8 64 |
| | | 3.68 | -20 8 72 |
| | 235 | 6.23 | 48 -40 -38 |
| | | 4.81 | 34 -42 -44 |
| | 460 | 5.94 | -42 -42 -42 |
| | | 5.77 | -34 -44 -40 |
| | | 4.16 | -40 -58 -42 |
| | 189 | 5.9 | -32 -40 -12 |
| | 1073 | 5.76 | -58 -60 10 |
| | | 5.22 | -22 -62 18 |
| | | 5.12 | -48 -64 4 |
| | 263 | 5.73 | -34 -86 32 |
| | | 4.96 | -38 -88 24 |
| | 387 | 5.57 | 60 -30 -18 |
| | | 5.2 | 58 -20 -14 |

| | | | |
|---|---|---|---|
| | | 4.16 | 50 -30 -14 |
| | 191 | 5.25 | 20 -58 16 |
| | 275 | 4.99 | -54 -10 26 |
| | | 4.82 | -46 -12 22 |
| | | 3.42 | -64 -10 18 |
| | 200 | 4.83 | -40 -10 -4 |
| | | 4.61 | -42 0 -2 |
| | | 3.55 | -48 -12 2 |
| | 209 | 4.53 | 30 54 -14 |
| | | 4.11 | 22 54 -16 |
| | | 3.79 | 22 36 -22 |
| | 173 | 3.99 | 18 58 22 |
| | | 3.7 | 30 56 14 |
| Semantic object > semantic event | 414 | 7.68 | -16 -86 -12 |
| | | 7.56 | -10 -90 -2 |
| | 607 | 6.97 | -28 -88 6 |
| | | 6.77 | -22 -94 14 |
| | | 6.77 | -24 -88 20 |
| | 578 | 6.88 | 18 -96 -2 |
| | | 6.67 | 12 -86 -2 |
| | | 6.29 | 36 -86 0 |
| | 128 | 6.67 | 28 -82 20 |

S7. ROI analyses from the response phase of the experiment: the AG showed significantly greater activation relative to all other conditions (all ts > 6.29, all ps < .000). The AG did not differ in response to the semantic-events and semantic-objects ($t(27) = 1.19$, $p = .25$), although both conditions differed to control ($t(27) = 3.40$, $p < .005$). With the exception of the autobiographical condition, all conditions showed significant deactivation relative to rest (all ts > -2.90, all ps < .005). The same pattern was found for the full DMN. Unlike the AG, the ATL showed significant great activation for all tasks relative to control (all ts > 6.22, all ps < .001), but no significant difference between conditions (all ts < 1.27, all ps > .22).
